# The effect of tissue composition on gene co-expression

**DOI:** 10.1101/492223

**Authors:** Yun Zhang, Jonavelle Cuerdo, Marc K. Halushka, Matthew N. McCall

## Abstract

Variable cellular composition of tissue samples represents a significant challenge for the interpretation of genomic profiling studies. Substantial effort has been devoted to modeling and adjusting for compositional differences when estimating differential expression between sample types. However, relatively little attention has been given to the effect of tissue composition on co-expression estimates. In this study, we illustrate the effect of variable cell type composition on correlation-based network estimation and provide a mathematical decomposition of the tissue-level correlation. We show that a class of deconvolution methods developed to separate tumor and stromal signatures can be applied to two component cell type mixtures. In simulated and real data, we identify conditions in which a deconvolution approach would be beneficial. Our results suggest that uncorrelated cell type specific markers are ideally suited to deconvolute both the expression and co-expression patterns of an individual cell type. Finally, we provide a Shiny application for users to interactively explore the effect of cell type composition on correlation-based co-expression estimation for any cell types of interest.

## Introduction

Cellular processes are governed by the interaction of genes and gene products. This produces shared patterns of expression between functionally related genes. Gene co-expression networks encode similarities in expression as edges in an undirected graph and have been used to identify genes that potentially share a regulatory relationship or common function (Butte et al., 2000; Basso et al., 2005; Voineagu et al., 2011; Yang et al., 2014).

### Co-expression network estimation

Statistical methods to estimate co-expression networks rely on gene expression measurements from biological replicates that share a common regulatory architecture. Relatively small variations in gene expression across these replicates are used to identify co-expressed genes. A major challenge is to design an experiment that is sufficiently large to distinguish true co-expression from noise while minimizing sources of technical variability, such as batch effects (Leek et al., 2010), that can easily overwhelm the biological signal (Parsana et al., 2017). As such, recent co-expression analyses have focused on data generated by large consortia such as the Gene Tissue Expression (GTEx) Project (Pierson et al., 2015; Saha et al., 2017).

The Pearson correlation coefficient is the most widely-used measure to capture linear dependencies between genes. For example, one of the most popular methods, WGCNA (Zhang et al., 2005; Langfelder & Horvath, 2008), begins by constructing a gene co-expression network based on the absolute value of the Pearson correlation coefficient. A challenge in the application of network estimation methods to gene expression data is that the number of genes is usually much greater than the number of samples. A variety of approaches, from the ad hoc to the statistically rigorous, have been used to address this challenge. For example, Schäfer & Strimmer (2005) proposed estimating the gene-gene covariance matrix by shrinking the empirical covariance matrix toward a diagonal matrix with unequal variance.

### Tissue composition and statistical deconvolution

Tissues are comprised of multiple cell types with distinct molecular phenotypes, resulting from cell type specific gene expression. Many of the cellular processes encoded by gene regulatory networks are specific to a particular cell type, resulting in fundamental differences in these networks across cell types.

A substantial number of statistical methods have been developed to estimate cellular composition and/or cell type specific RNA expression from tissue-level expression data. The vast majority of these methods are based on a linear combination of cell type specific expression profiles, originally proposed by Venet et al. (2001). Specifically, these methods model the observed expression, *Y_ij_*, of gene *i* in tissue sample *j* as a linear combination of cell type specific expression, *X_ik_*, of each of the *K* cell types that make up the tissue:

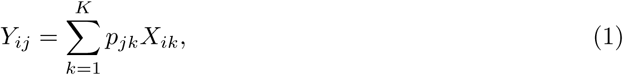
where the compositional proportions have the constraint 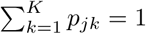. A clear limitation to this model is that cell type expression is assumed to be constant across tissue samples, *X_ik_* is the same for all *j*. This is biologically implausible and often in direct opposition to subsequent analyses that seek to identify genes whose expression differs between groups of samples.

Two recent methods, focusing on tumor/normal mixtures, have improved upon these earlier approaches by allowing cell type expression to vary across samples. One method, ISOpure, assumes that each normal expression profile is a convex combination of a set of reference normal profiles and that the cancer profiles are similar to one another (Anghel et al., 2015). The other method, DeMix, provides flexible modeling of mixed tissue samples across four different situations: with or without reference gene profiles and with or without matched tumor/normal samples. In each of these situations, DeMix assumes that the normal mixture component can be measured directly (Ahn et al., 2013).

In this study, we begin by illustrating the challenges of estimating co-expression in the constituent cell types from heterogeneous tissue samples. We then assess the ability of deconvolution methods to facilitate the estimation of cell type specific co-expression. Finally, we conclude with a discussion of the conditions in which deconvolution improves estimation of cell type specific co-expression. For the purpose of this study, we focus on correlation-based co-expression networks, which are arguably the most commonly used (Villaverde & Banga, 2014; Petereit et al., 2016) and have been shown to perform comparably with more complex methods (Song et al., 2012).

## Methods

### Co-expression network estimation

In the subsequent analyses, we use the absolute value of either Pearson’s correlation coefficient or a shrinkage correlation estimator (Schäfer & Strimmer, 2005) to assess the pairwise association between genes. The latter is a modification of Pearson’s correlation coefficient with a shrinkage intensity parameter, λ ∈ [0,1]. Before constructing the shrinkage estimator, we define notation for the empirical variance-covariance matrix, 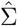. Suppose 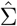 is a *m* × *m* real, symmetric, and positive definite matrix, then it can be expressed as:

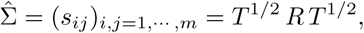
where

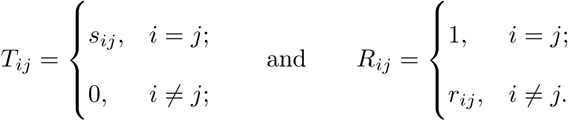

In other words, the off-diagonal elements (*i* ≠ *j*) have the following expression:

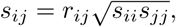
where *r_ij_* is Pearson’s correlation coefficient.

The modified correlation estimator is a linear shrinkage approach that combines the diagonal matrix with unequal variances, *T*, and the empirical variance-covariance matrix, 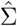, such that

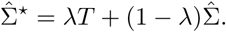

In the above equation, *T* is the shrinkage target, and the optimal shrinkage parameter is

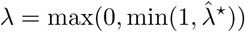
with

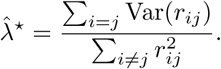

This optimal shrinkage parameter is determined *analytically* based on the minimal mean squared error criterion, which is derived by Ledoit & Wolf (2003). Finally, the shrinkage estimators can be written explicitly as

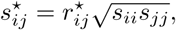
where 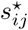 it the (*i*, *j*)*th* element of 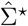, and 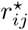 is the corresponding shrinkage correlation estimate. This approach is implemented in the R package `GeneNet`.

### Sample-specific deconvolution

A *sample-specific* deconvolution model can be expressed as:

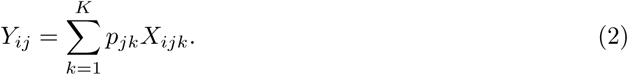

Note that Equation (2) is a generalization of Equation (1) where cell type specific expression *X_ijk_* is now allowed to vary across the sample index *j*. Note that co-expression analysis for a specific cell type is now possible because the expression within each cell type is allowed to vary across samples.

However, in the most general case where only the *Y_ij_* are observed, Equation (2) is an *under-determined* system of equations, i.e. the number of equations is less than the number of unknown parameters. With further assumptions, the deconvolution problem can be categorized into two types: (i) *full* deconvolution for unknown *p_jk_* and *X_ijk_*, and (ii) *partial* deconvolution with known *p_jk_* or *X_ijk_* (Mohammadi et al., 2017).

### ISOpure deconvolution

ISOpure (Quon et al., 2013; Anghel et al., 2015) was originally developed for tumor profiles (pre-purification) that are considered as mixtures of cancer and normal profiles. By comparing the tumor profiles to a set of unmatched normal profiles, ISOpure estimates tumor purities (i.e. proportions in the mixtures) and individual cancer profiles (post-purification) for each tumor sample. The ISOpure statistical model is based on a Dirichlet-multinomial mixture model. The ISOpure algorithm estimates the proportions and the purified expression profiles in two steps. The first step is a Bayesian approach that iteratively updates the Dirichlet prior for the proportions and the compound multinomial distribution for the mixed profiles after appropriate data reorganization. Based on the outputs from the Bayesian model, the second step is to estimate the individual profiles for the target cell type by maximum likelihood.

For a mixture of two cell types, we adopt ISOpure to decompose the compositional cell type profiles using the R package `ISOpureR`. For the two cell types, suppose cell type 1 is the target cell type, whose profile is in lieu of the cancer profile, and cell type 2 is the reference cell type, whose profile is in lieu of the normal profile. ISOpure assumes that there is a set of reference cell type profiles that can represent the cell type 2 expression profile. Thus, ISOpure inputs *Y_ij_* (i.e. the mixed profiles) and *X_ij_*_2_ (i.e. the reference profiles), where *j*’s from the two collections of profiles do not need to match; and, it returns estimated *p_j_*_1_ and *X_ij_*_1_, where the *j*’s correspond to the mixed profile indices. Normalized but *not* log-transformed data are require for ISOpure. In our analyses, we use quantile normalization (Bolstad et al., 2003) to make expression profiles comparable from sample to sample. Technically, ISOpure depends on an initial random seed, and the algorithm may fail to converge occasionally. When this happened, we repeated the analysis with a new random seed.

### Limitations of regression-based composition adjustment

The full deconvolution problem is an under-determined system of linear equations, even in the case of a mixture of two cell types. Sometimes, investigators may have knowledge of *p_jk_* through complementary measurements (e.g. histological imaging). In this case, a regression-based deconvolution model (Glass & Dozmorov, 2016) has been proposed such that

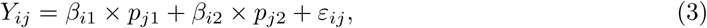
where *β_i_*_1_ and *β_i_*_2_ are the unknown average theoretical cell type-specific gene expression and *ε_ij_* represents random error. This model suffers from two related limitations. First, *ε_ij_* captures technical variation but not biological variation between samples. Second, as previously shown in Jaffe & Irizarry (2014), Equation (3) is only valid when the difference between gene expression in each sample and the cell type average are the same across cell types, which is rarely a reasonable assumption.

### Simulations

We utilize simulations to access the effects of cell type mixture and deconvolution on co-expression networks. Let *m* be the number of features, i.e. *i* = 1,…, *m*. Using vector forms, we denote **Y***_j_* = (*Y*_1_*_j_*,… *,Y_mj_*)′ for the *j*th mixed expression profile, and **X**_jk_ = (*X*_1_*_jk_*,…, *X_mjk_*)′ for the pure cell type-*k* expression profile. Also, we denote the log2-transformed profiles as 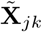 and 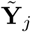, such that:

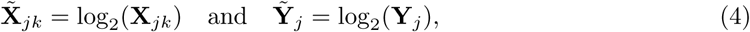
and similarly for the mixed profiles. We emphasize this log-transformation because it has been shown that deconvolution models (Equation (1) or Equation (2)) should be built upon data that have not been log-transformed, since such a transformation introduces bias in the resulting profiles (Zhong & Liu, 2012).

In our simulations, we generate the log-transformed data, 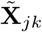, from an m-dimensional multivariate normal distribution, and use Equation (4) to obtain the appropriate data for deconvolution, **X***_jk_*. Focusing on two cell types, we randomly draw 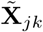 from the following multivariate normal distribution

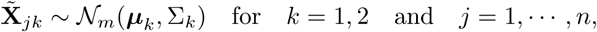
where *n* is the number of samples for each cell type. Suppose the cell type specific mean vectors, ***μ****_k_*, are available. We design the cell type specific covariance matrix (i.e. co-expression structure), Σ*_k_*, according to Figure 1. For the two cell types, we set one as the reference cell type with an identity covariance matrix and the other cell type as the target with a block-diagonal covariance matrix. The block-diagonal covariance matrix is further characterized by the number of blocks, block size, correlation magnitude, and co-varying features: the most differentially expressed, randomly selected, or the least differentially expressed.

**Figure 1:**
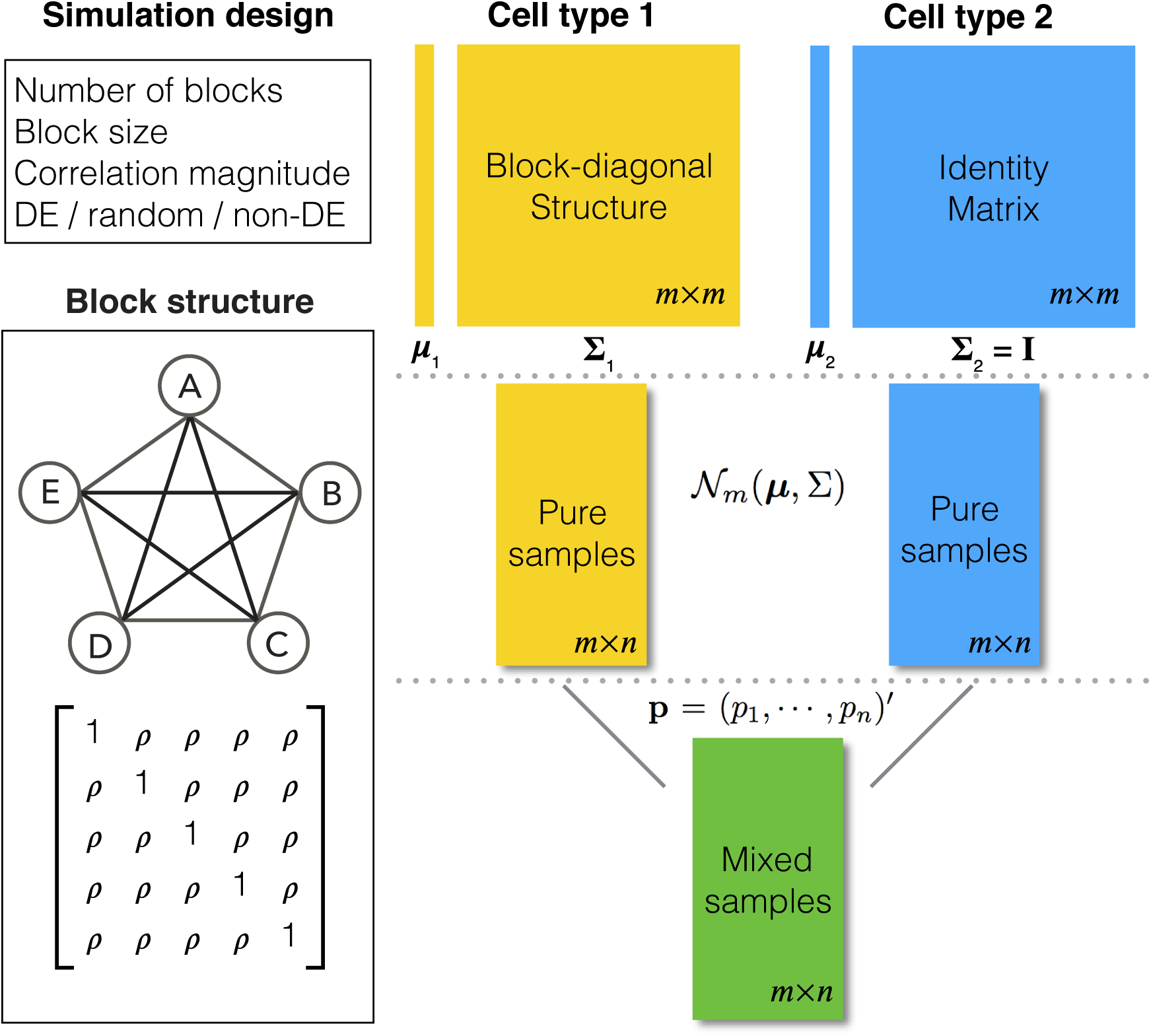
Overview of the simulation scheme. Consider two cell types: cell type 1 (the target cell type) in yellow with known mean vector and block-diagonal covariance matrix, Σ_1_, and cell type 2 (the reference cell type) in blue with known mean vector and identity covariance matrix, Σ_2_. **The top-left panel** characterizes the design of Σ_1_. The co-expression signal in the target cell type is concentrated on the most differentially expressed (DE) features, randomly selected features, or the least differentially expressed (non-DE) features. **The bottom-left panel** is an example of the type of structure (i.e. co-expression network) applied to each signal-receiving block, where A – E are five genes, and the matrix beneath specifies the covariance structure. **The right panel** is a flow chart of the data generation procedure. Pure samples are generated from a multivariate normal distribution, 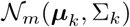; mixed samples are mixtures of the pure samples in given proportions, **p** for the target cell type and **1** − **p** for the reference cell type.

With ***μ***_1_, ***μ***_2_, Σ_1_, Σ_2_ specified, we generate random samples for pure cell type 1, 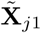, and pure cell type 2, 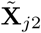, using the corresponding multivariate normal distributions. Before mixing, we transform the data to obtain **X***_j_*_1_ and **X***_j_*_2_. Let **p** = (*p*_1_,…, *p_n_*)′ be the proportions of cell type 1 and 1 − **p** be the proportions of cell type 2 in the mixture. Finally, expression in the mixed samples is generated as follows:

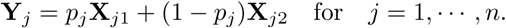

### The microRNAome data

We obtained cell type-specific microRNA expression data from the Bioconductor package `microRNAome` (McCall et al., 2017). Simulated data were generated according to Figure 1 using 382 well-characterized microRNAs in pure Aortic Smooth Muscle Cells (ASM) and pure Aortic Endothelial Cells (AEC). Without loss of generality, we set AEC as the reference cell type. For each cell type, we generated 20 samples and computationally mixed these to obtain 20 mixed samples. We designed the co-expression pattern for ASM as follows: three co-expression blocks, 10 features per block, and a correlation magnitude of 0.7. The 20 mixture proportions were equally spaced from 0 to 1. In different simulations, the co-expressed features were selected as the most differentially expressed, as the least differentially expressed, or randomly.

We applied the ISOpure deconvolution algorithm to the mixed samples and compared the simulation performance using ROC curves based on the Pearson correlation and the shrinkage correlation for the pure ASM, mixed, and deconvoluted samples.

To assess performance on a real data network, we selected two cell types from the microRNAome data: dendritic cells (18 samples) and fat cells (15 samples). We set the fat cells as the reference cell type. We identified 363 microRNAs, which were non-zero in at least half of the samples for each cell type. We mixed the pure cell type samples with two different sequences of mixture proportions. In terms of the proportions of the dendritic cells, the first sequence is equally spaced from 0 to 1, and the second sequence is equally spaced from 0.4 to 0.6. Based on Equation (2), we obtained computationally mixed samples by randomly selecting 15 samples of pure dendritic cells and combining them with the 15 samples of pure fat cells in the given proportions. This mixed sample generation was repeated for 20 times to produce 20 complete mixed data sets.

A true edge in the real data network was defined based on the empirical correlation matrix from the data. We dichotomize the empirical correlation matrix and obtain the edge matrix as follows:

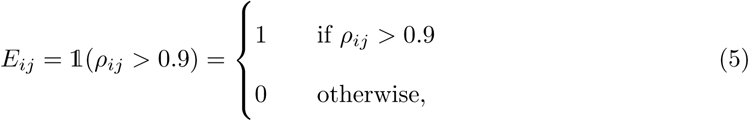
where *E_ij_* = 1 indicates an edge between the *i*th and *j*th features, *E_ij_* = 0 indicates no edge, 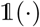 is an indicator function, and *ρ_ij_* is the Pearson correlation between the pair of features.

### Assessments of performance

We used receiver operating characteristic (ROC) curves (Hanley & McNeil, 1982) and area under curve (AUC) statistics to assess network estimation based on correlation measures. The ROC curve shows the relationship between the true positive rate (TPR) and the false positive rate (FPR) for a range of correlation thresholds, where:

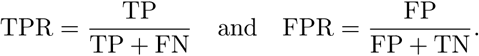

At each threshold, correlation values above the threshold are considered as positives (i.e. edges in a network), and correlation values below the threshold are negatives (i.e. no edge). In the simulations, true edges are defined as any non-zero off-diagonal element in Σ_1_.

An ROC curve close to the top-left corner of the plotted area indicates high TPR and low FPR. On the contrary, if a curve is close to the diagonal line, it suggests that the performance is roughly the same as random chance. The AUC statistic is a quantitative summary of the ROC curve, which ranges from 0 to 1. As the ROC curve approaches the top-left corner, the corresponding AUC statistic approaches 1.

## Results

### An illustrative model of tissue level gene expression

Consider the following model of tissue level gene expression arising from a mixture of two cell types:

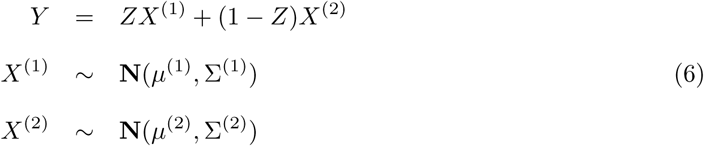
where *Y* is a vector of tissue-level gene expression, *X*^(1)^ and *X*^(2)^ are random vectors of gene expression in cell types 1 and 2 respectively, and *Z* is a scalar random variable denoting the proportion of cell type 1 in the tissue. In contrast to a Gaussian mixture model, in which each observation comes from only one of the mixture components, i.e. *Z* ∈ {0, 1}, here each observation is a convex combination of cell type specific expression vectors, i.e. *Z* ∈ (0,1). Note that conditional on the mixing proportion,

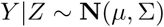
where

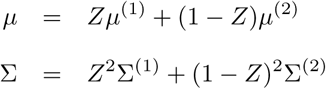

Thus, the covariance between any two genes *Y*_1_ and *Y*_2_ can be expressed as:

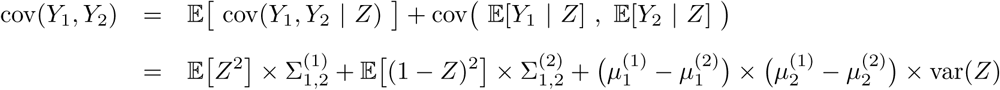

If tissue composition is variable, i.e. var(*Z*) > 0, the covariance between genes in the mixed tissue depends on the covariances in each cell type, as well as the difference in expression between the cell types. Finally, the correlation between any two genes *Y*_1_ and *Y*_2_ can be expressed as:

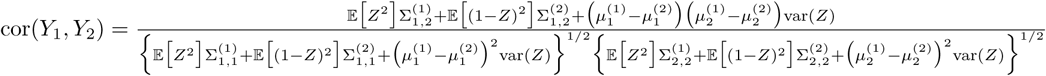

Note that even if the genes are positively correlated in both cell types, their tissue-level correlation can be negative if the differences in average expression between cell types for the two genes differ in sign and the variance in the mixing proportion is sufficiently large.

To further examine the effect of variable composition on tissue-level gene expression correlation, consider the following simplifications:

- 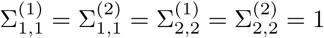: the cell type variances are all equal to 1
- 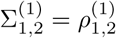: the two genes are correlated in cell type 1 and have correlation magnitude equal to some non-zero 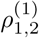
- 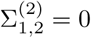: the two genes are not correlated in cell type 2
- 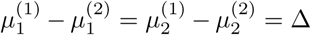: the differences in expression between cell types are equal to Δ

In this case, the tissue-level correlation can be expressed as:

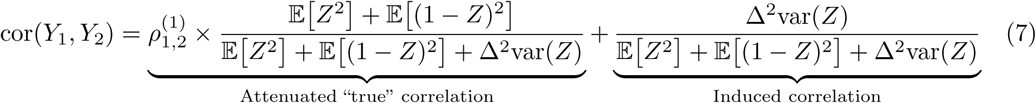

The first part of the summation represents the attenuated correlation due to the cell type mixture. The second part of the summation represents the correlation induced by variation in the mixing proportion. To be noted, the induced correlation also depends on how differently these genes are expressed in the different cell types.

Figure 2 provides an illustrative example of between-gene correlation induced by variable mixing of cell types in tissue samples. In this simulated experiment, we consider 10 genes (*a–j*) measured in two cell types (100 samples per cell type), where five of the genes are differentially expressed between the two cell types (Figure 2A). As in Equation (6), we assume that gene expression in each cell type follows a multivariate normal distribution. For illustrative purposes, we assume that Σ^(1)^ is block diagonal and Σ^(2)^ is the identity matrix (Figure 2B–C). We set *ρ*^(1)^ = 0.7 for the correlated gene pairs, *μ*^(1)^ − *μ*^(2)^ = 5 for the differentially expressed genes, and var(*Z*) = 0.03. Between the three mixtures (Figure 2D–F), we only vary the expected value of the mixing proportion, 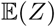. Specifically, we consider generating the mixing proportions from three beta distributions (Figure 2G–H). As expected, in the mixed tissue samples, one can clearly see the effects of both the cell type specific correlation and the correlation induced by the mixing of cell types (comparison between Figure 2B–C and Figure 2D–F). For genes *g–i*, which are not differentially expressed between the two cell types, we observe a clear attenuation of the correlation structure as the proportion of cell type 1 decreases (moving from Figure 2D to Figure 2F). For genes *a–e*, which are differentially expressed between the two cell types, the correlation induced by the mixture masks the cell type specific correlation even in an equal mixture of the two cell types (Figure 2E). Finally, the amount of correlation induced by the mixture also depends on the statistical properties of the mixing distribution. Specifically, it is a non-monotone function of the first two moments of the distribution of the mixing proportion.

**Figure 2:**
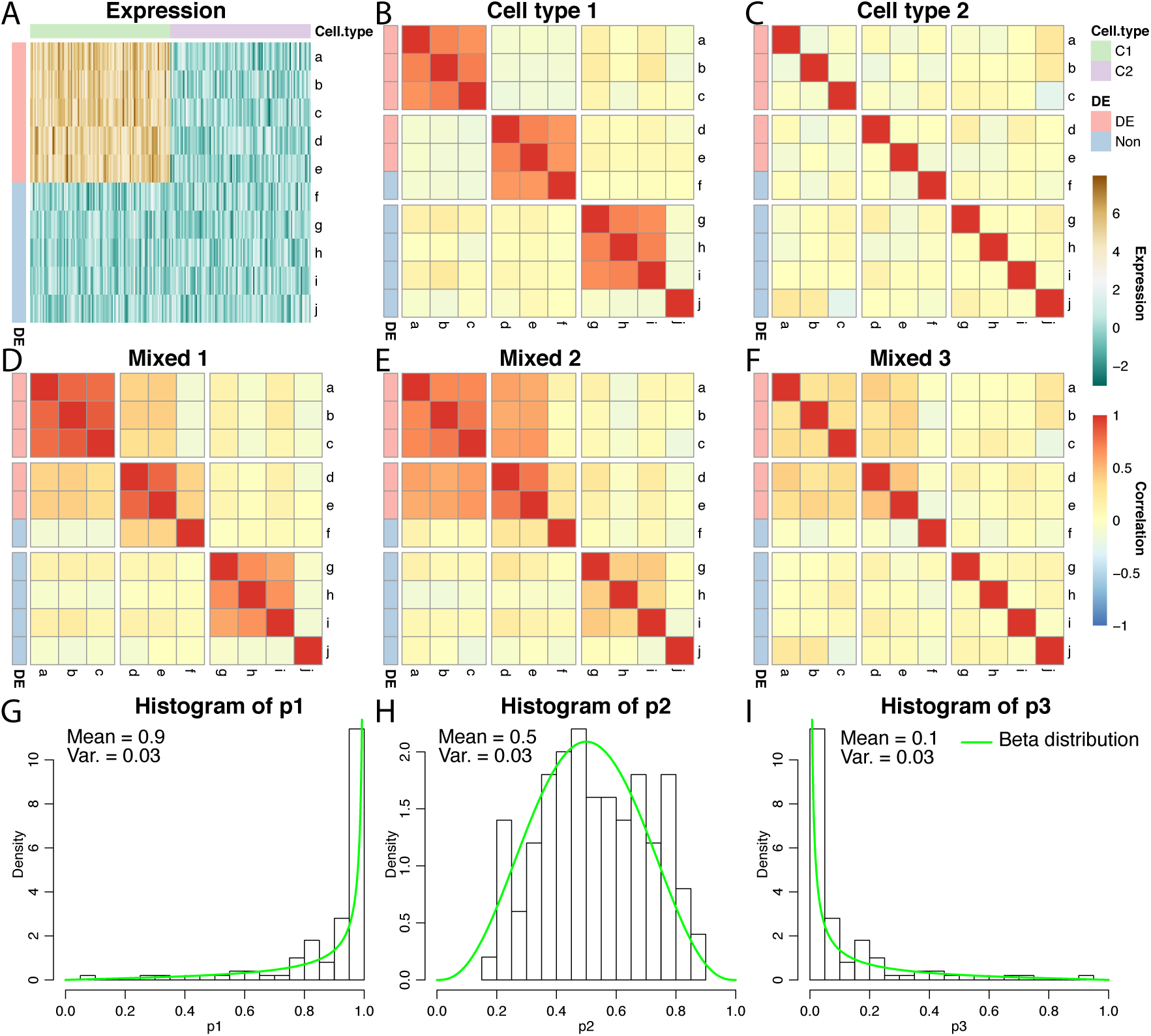
An example of correlation induced by cell type mixture. Consider 10 hypothetical genes (namely, *a–j*) that are expressed in two cell types. Expression profiles of 100 samples for each cell type are plotted in panel A. The first five genes (*a–e*) are differentially expressed between the two cell types, and the other five genes (*f–j*) are not differentially expressed. Panels B–F are heatmaps of correlations. These genes form three co-expression blocks in cell type 1 (panel B) and are not co-expressed in cell type 2 (panel C). We generated three mixtures (panels D–F) with varying proportions of cell type 1 drawn from beta distributions (panels G–I).

### Co-expression estimation in pure, mixed, and deconvoluted samples

Figure 3 shows the performance of two correlation-based network estimators on simulated data from pure cell type samples, samples from a variable mixture of two cell types, and the corresponding deconvoluted samples, with three types of co-expressed features. Both estimators performed best on the pure cell type data (AUC ≥ 0.94), which suggests that both correlation measures are able to capture the co-expression signal well in pure samples. Compared to the pure samples, the mixed samples largely deviate from the true signals and have poor performance. Figure 3A shows the results when only the most differentially expressed features are co-expressed. Using the Pearson correlation, the mixed ROC curve is close to the pure curve. This is because the Pearson correlation is increased due to the correlation induced by the cell type mixture, which does not represent cell type specific co-expression. In contrast, the shrinkage correlation estimator (Figure 3D) is a more conservative measure, which regularizes the empirical covariance matrix towards a diagonal matrix. This appear to greatly reduce the effect of the induced correlation. Finally, the deconvolution produces modest to substantial improvement over the mixed samples, except in the situation noted above. Across all scenarios, the deconvoluted samples result in stable performance (AUC > 0.8).

**Figure 3:**
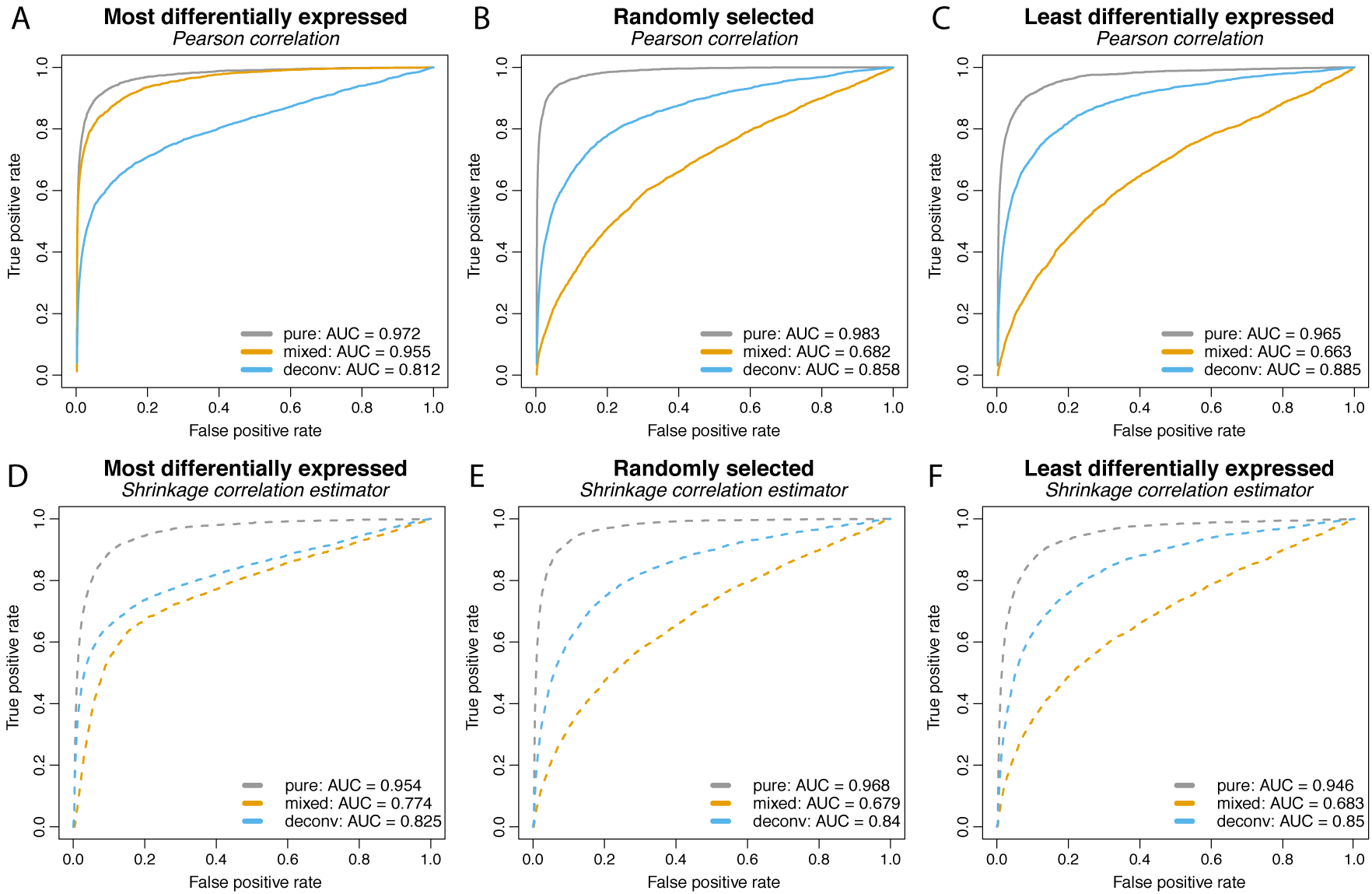
ROC curves from a simulation study with number of blocks = 3, block size = 10, number of samples = 20, and correlation magnitude = 0.7. The mixture proportions of cell type 1 were equally spaced from 0 to 1. Block correlation structure was imposed on the most differentially expressed features (left column), randomly selected features (middle column), and the least differentially expressed features (right column). Differential expression was quantified by the absolute difference between the cell type-specific mean vectors. Two measures of co-expression were used: the Pearson correlation (top row) and the shrinkage correlation implemented by `GeneNet` (bottom row). In each panel, the ROC curves for the pure samples (gray curve), the mixed samples (yellow curve), and the ISOpure deconvoluted samples (blue curve) are shown.

To further evaluate the deconvolution performance, we plotted the estimated proportions against the true proportions for the target cell type in Supplementary Figure S1. Recall that the true proportions in this simulation are equally spaced from 0 to 1. Based on the error measures, mean absolute difference (mAD) and the root mean squared distance (RMSD) (Mohammadi et al., 2017), the deconvolution estimation of mixing proportions appears stable and robust to the choice of co-expression features.

Additionally, we conducted simulation studies with mixing proportions randomly generated from a beta distribution. We show results similar to Figure 3 for these simulations in Supplementary Figures S2, S3 and S4. Overall, they reveal a similar pattern as that seen in Figure 3. The performance of the deconvoluted curves are stable across different scenarios and robust to the selection of co-expression features. They effectively recover the true co-expression signal from cell type mixtures. Without deconvolution, the mixed curves vary dramatically from case to case. When only a small proportion of the target cell type is present in the mixture, the mixed curve is close to the diagonal line (Figure S2); when a large proportion of the target cell type is present in the mixture, the mixed curve performs similarly to the deconvoluted curve (Figure S4).

### Appropriate cases for deconvolution

In order to further evaluate the situations in which deconvolution is able to recover the pure sample co-expression, we generated more simulation scenarios by varying block size, number of samples, and correlation magnitude in the same simulation design. While one parameter is varied, other parameters are kept fixed as before. The resulting AUC values are shown in Figure 4.

**Figure 4:**
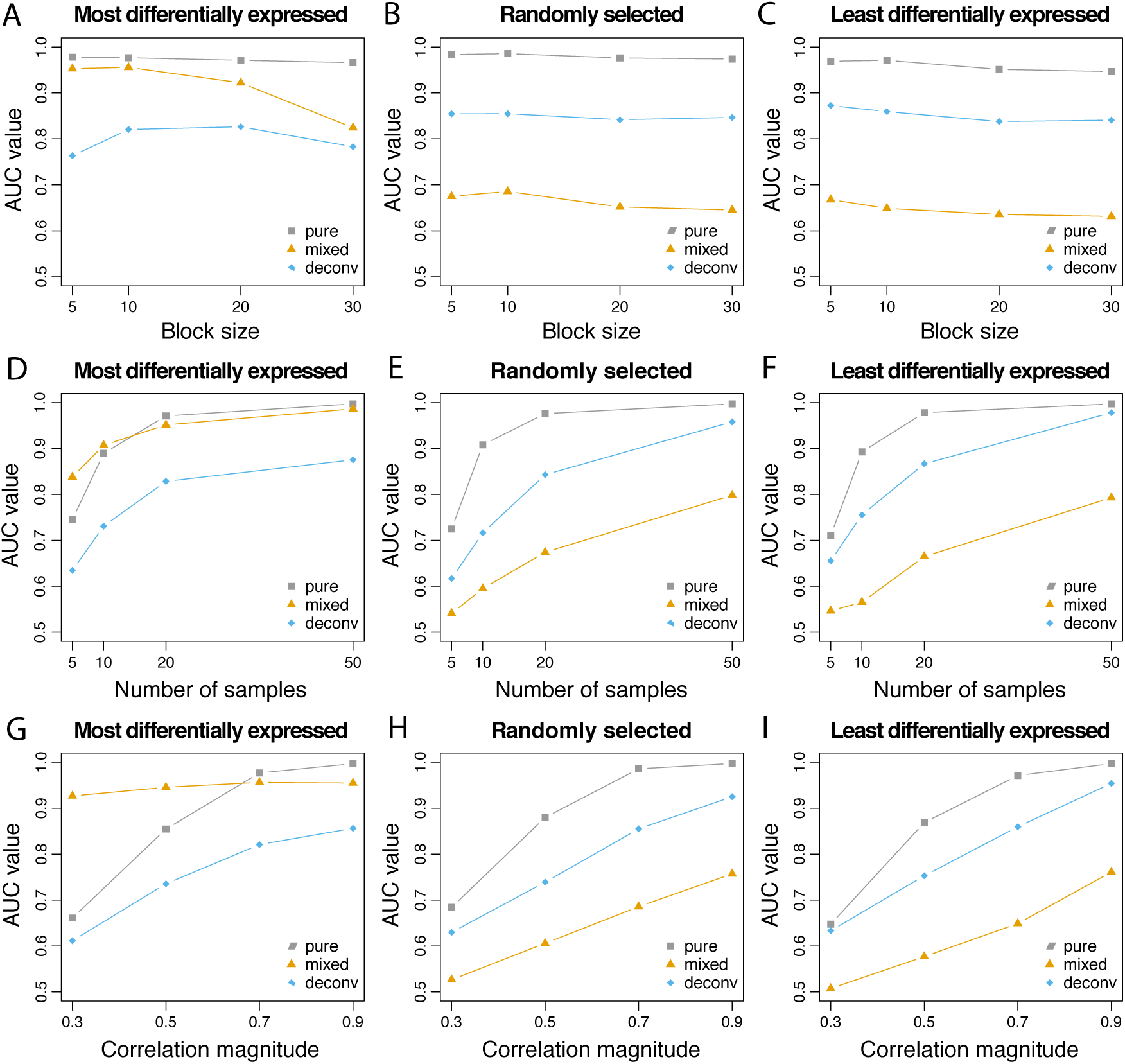
AUC values are reported with varying block size, number of samples, and correlation magnitude. Co-expression association is measured by Pearson correlation. When one parameter is varied, the others are fixed at: block size = 10, number of samples = 20 and correlation magnitude = 0.7. As before, three correlated blocks are generated.

In the first column of Figure 4, in which the most differentially expressed genes are also those that are co-expressed, the AUC values for the mixed samples are higher than those for the deconvoluted samples, and sometimes even higher than that of the pure samples. As shown in Equation (7), if the correlated features are also differentially expressed between the two cell types, the differential expression (Δ) induces correlation in the mixture if the mixture proportion varies across samples. The high AUC values are attributable to the induced correlation. When the true correlations are difficult to estimate due to small sample size (Figure 4D) or of small magnitude (Figure 4G), the induced correlation dominates the attentuated co-expression signal in the mixture. This results in the mixed samples having higher AUC than the pure samples.

In the second and third columns of Figure 4, in which the co-expressed features are randomly selected or the least differentially expressed, the AUC values are always higher for the pure samples than the deconvoluted samples, which in turn are higher than the mixed samples. Overall, the AUC values increase as the number of samples or the correlation magnitude increases. The block size does not appear to impact the co-expression network estimation performance.

In summary, the ISOpure deconvolution method recovers the *true* co-expression signal from mixed samples. If the co-expressed features are differentially expressed between the two cell types, the induced correlation may lead to better estimation of the co-expression network; however, caution is required in interpreting the result as these features would appear correlated in the mixture regardless of whether they were truly co-expressed in the target cell type.

### Real data networks

Using the `microRNAome` dataset, we computationally mixed pure dendritic cells with the same number of pure fat cells as described in the Methods. Treating the empirical network in the pure cell type data as the true network, we plot the ROC curves for network reconstruction with the mixed and deconvoluted profiles (Figure 5B,D) with respect to two mixing proportion schemes. In both ROC plots, we focus on the region where the false positive rate is smaller than 0.5, and observe that the deconvoluted curve is uniformly better than the mixed curve. Over the whole range of false positive rate, the AUC statistics are also slightly higher for the deconvoluted curve in both cases (mixed vs. deconv.: 0.711 vs. 0.723 (Figure 5B), 0.824 vs. 0.845 (Figure 5D)). Again, the real data confirms the improvements gained from deconvolution. For both sequences of mixing proportions, estimation of very small and very large proportions show less variation (Figure 5A,C). The gain from deconvolution is larger when there is greater variation in the mixture proportions (Figure 5B).

**Figure 5:**
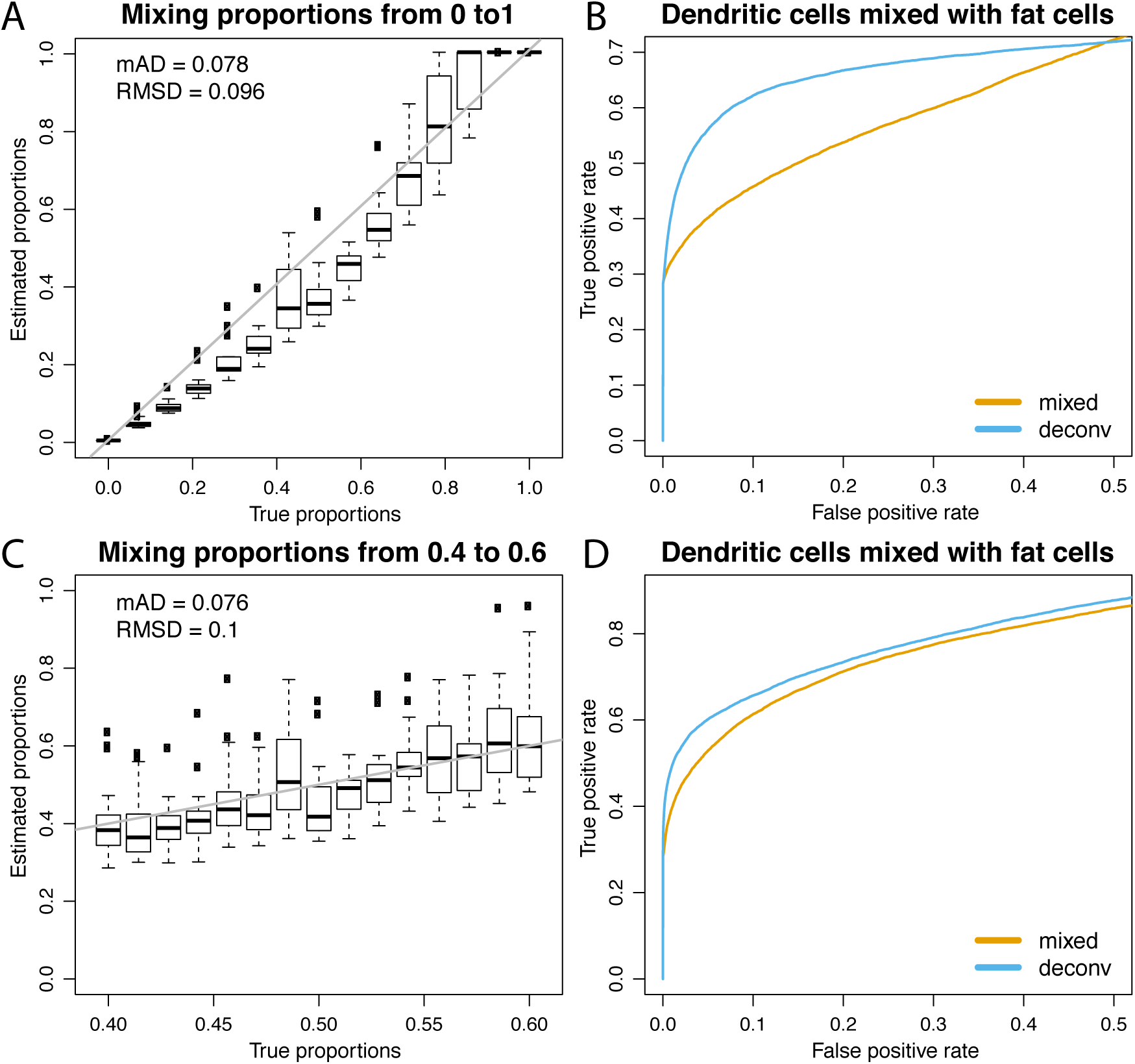
Deconvolution performance on real data. Top row: mixing proportions varying from 0 to 1. Bottom row: mixing proportions varying from 0.4 to 0.6. Panels A and C show the estimated proportions versus the true proportions; the gray line is the reference line for both proportions being equal. By defining a true edge if absolute correlation > 0.9, panels B and D show dominating ROC curves for the deconvoluted samples over the mixed samples in the zoomed region where false positive rates < 0.5.

Lastly, we varied the threshold used to define the true network based on the empirical correlation matrix. Supplementary Figure S5 shows the corresponding AUC values for varying thresholds. As the threshold increases, we observe an increase in the AUC due to a stronger signal defined in the empirical network. The deconvolution AUC remains uniformly better than the mixture AUC.

### An example of co-expressed microRNAs

Using real data, we illustrate what would happen to a pair of co-expressed microRNAs when mixing two cell types and deconvoluting the mixed samples. Figure 6 shows scatter plots of two microRNAs, *hsa-let-7a-5p/7c-5p* and *hsa-miR-766-3p*, in two different cell types. In dendritic cells (Figure 6A), these two features are strongly positively correlation (cor ≈ 0.9); in fat cells (Figure 6B), these two features appear uncorrelated (cor ≈ 0). After computationally mixing the two cell types with mixing proportions ranging from zero to one. The mixed samples (Figure 6C) appear negatively correlated (cor ≈ −0.3), which is entirely induced by the mixing. Finally, after ISOpure deconvolution targeting expression in the dendritic cells (Figure 6D), the positive correlation (cor ≈ 0.9) is re-established between the same pair of features.

**Figure 6:**
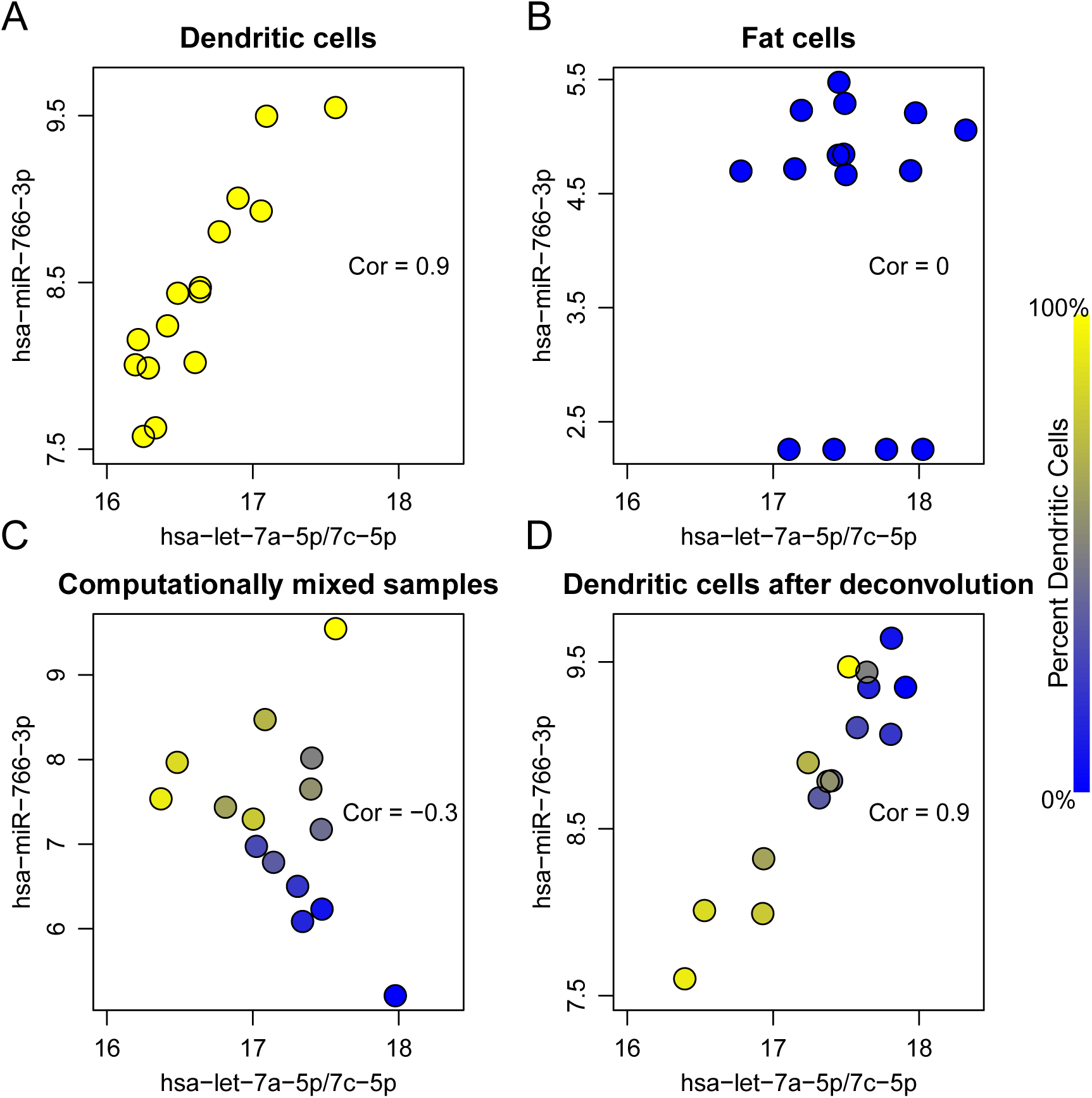
A real data example of two microRNAs (represented in the axes), which are highly correlated in dendritic cells (panel A) but uncorrelated in fat cells (panel B). Computationally mixed samples do not preserve the correlation (panel C). After deconvolution, strong correlation is recovered in the estimated expression of dendritic cells (panel D). Correlation values are shown in corresponding scatter plots. The color of each dot represents the proportion of dendritic cells in that sample.

### Interactive exploration of correlation induced by tissue composition

To further explore tissue composition induced co-expression, we developed a Shiny application that allows the user to easily evaluate the degree of induced or attenuated correlation due to cell type mixtures. The app allows users to select values for the parameters indicated in Figure 1. The user must first select a dataset to work with (currently, the app is tailored to work with two datasets: celltype and microRNAome), which will automatically update the list of cell types to choose from in the two dropdown menus for selecting cell types. The user can then proceed to select other parameters for the simulation. We have also added the option to generate beta-distributed proportions, for which the user can specify the mean and variance. Once all parameters have been selected, the user can generate a ROC curve to compare performance based on mixed and pure samples.

## Discussion

In the absence of targeted perturbation experiments, measurement of co-expression between genes is the primary method of assessing gene-gene interactions. The Pearson correlation between genes is usually the first step in co-expression network reconstruction, such as the widely-used WGCNA algorithm. By using this fundamental measure, our results are applicable to a wide range of network algorithms based on the Pearson correlation.

As we have shown, gene co-expression in tissue samples is often dominated by varying cellular composition. When applied to tissue gene expression data, methods that rely on co-expression to define gene modules, primarily identify groups of genes specific to a given component cell type. Variation in these gene modules therefore is capturing changing tissue composition. Some of these compositional differences may arise from biological variation or disease processes that affect the entirety of the tissue from which the sample was obtained. However, others may be due to spatial heterogeneity within a tissue, such that the proportion of cell types within the tissue sample is not representative of the proportion of cell types within the entire tissue. For example, variable sampling of highly localized cellular structures within in a tissue can result in substantial variation in tissue sample composition (McCall et al., 2016). These latter sources of co-expression reflect technical rather than biological variation.

In a mixture of two cell types, current deconvolution methods can be used to estimate cell type specific expression within each sample. Their ability to provide accurate estimates of gene-gene correlation depends upon the covariance between marker genes within each component cell type. When the genes used to identify the abundance of a specific cell type are all highly correlated with each other (e.g. members of a cell type specific pathway), current methods are unable to distinguish between changes in composition and changes in expression. Therefore, ideal marker genes are those that are cell type specific but uncorrelated within each cell type.

In this manuscript we have focused on correlation-based network estimation; however, two other co-expression network estimation categories should be noted: information-theoretic and Bayesian (Villaverde & Banga, 2014). Mutual information (MI) is an information-theoretic measure that captures *nonlinear* dependences between genes. Another popular network estimation method, ARACNE (Basso et al., 2005; Margolin et al., 2006), begins by calculating the mutual information between each pair of genes. Finally, Bayesian networks encode causal relationships or hierarchical structure between genes in a *directed acyclic graph*. A rich collection of Bayesian network learning algorithms are implemented in the R package `bnlearn` (Scutari, 2009).

An alternative to analyzing complex tissue samples is to measure cellular expression in a more homogeneous population obtained from cell culture, laser capture microdissection, centrifugation, or fluorescence activated cell sorting (FACS). These methods simplify the assessment of cell type specific co-expression; however, they often fail to determine the true biology of an organ where cell-cell interactions are critical to transcriptomic expression. Moreover, these methods often result in residual compositional heterogeneity, the introduction of technical artifacts, expression changes due to cell culture, and/or RNA degradation (Debey et al., 2004; Shen-Orr & Gaujoux, 2013). Therefore, it is often necessary to estimate cell type specific co-expression from tissue gene expression data.

## Supplementary Figures

**Figure S1:**
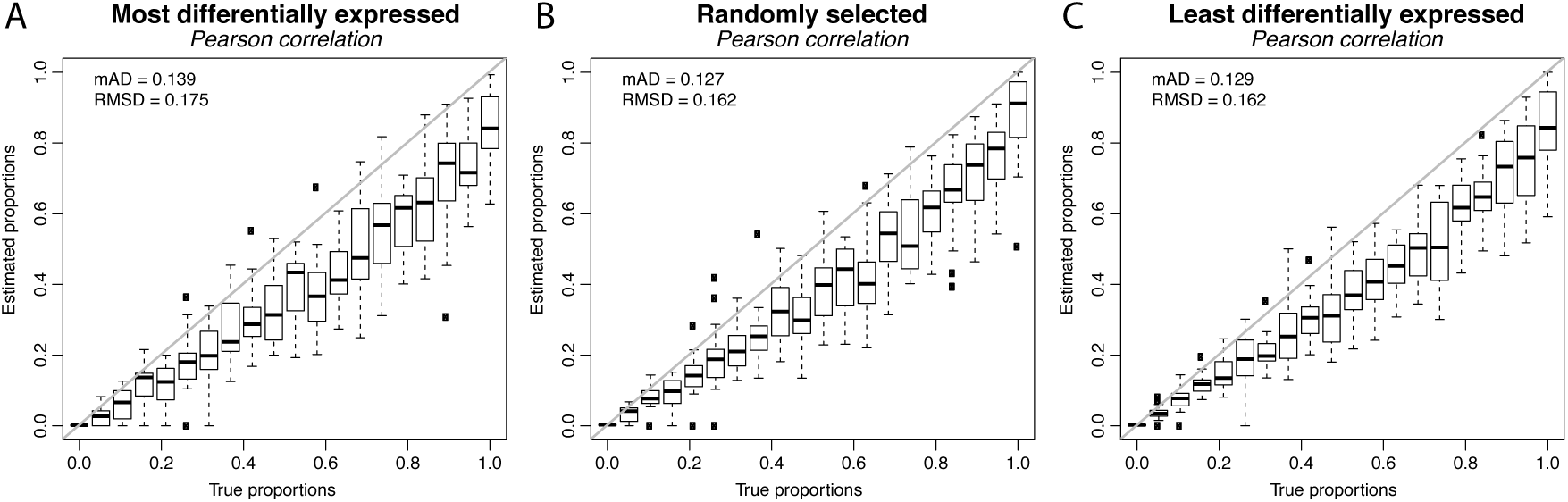
Distribution of estimated proportions corresponding to the simulations in Figure 3.

**Figure S2:**
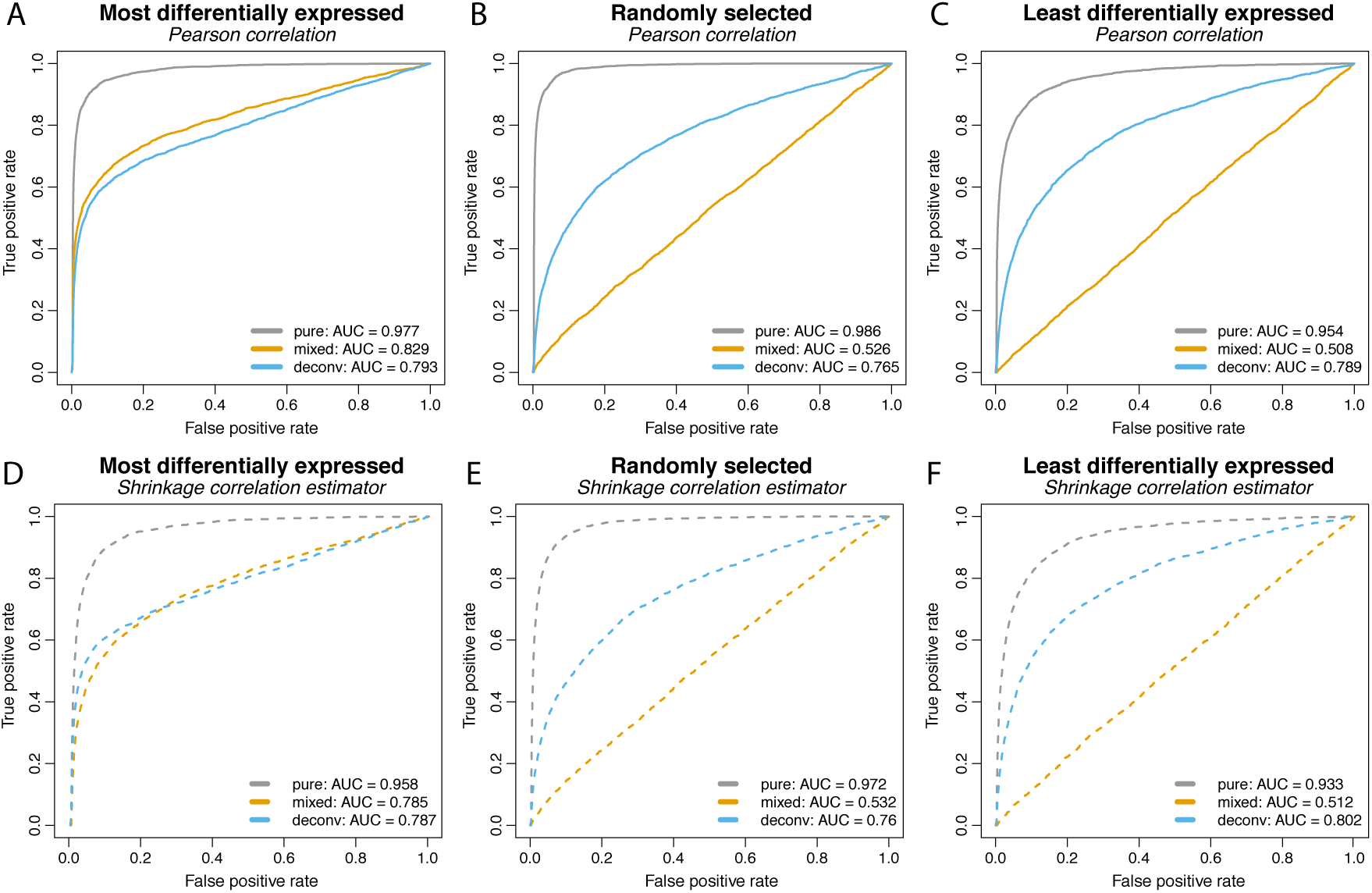
Counterpart of Figure 3 with proportions generated from the Beta(2,8), i.e. small proportions (on average 0.2) for signal-carrying cell-type in the mixture.

**Figure S3:**
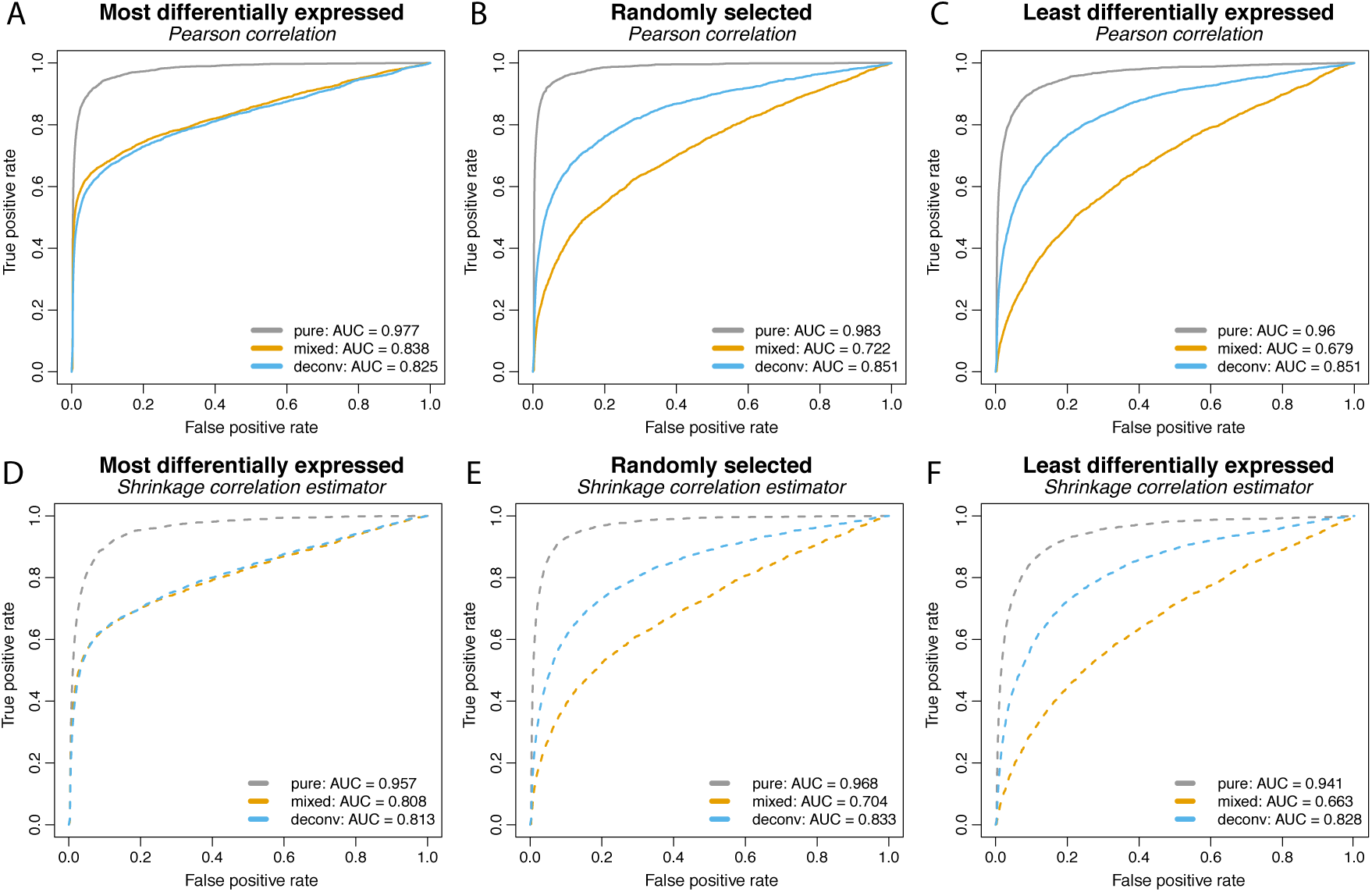
Counterpart of Figure 3 with proportions generated from the Beta(5,5), i.e. median proportions (on average 0.5) for signal-carrying cell-type in the mixture.

**Figure S4:**
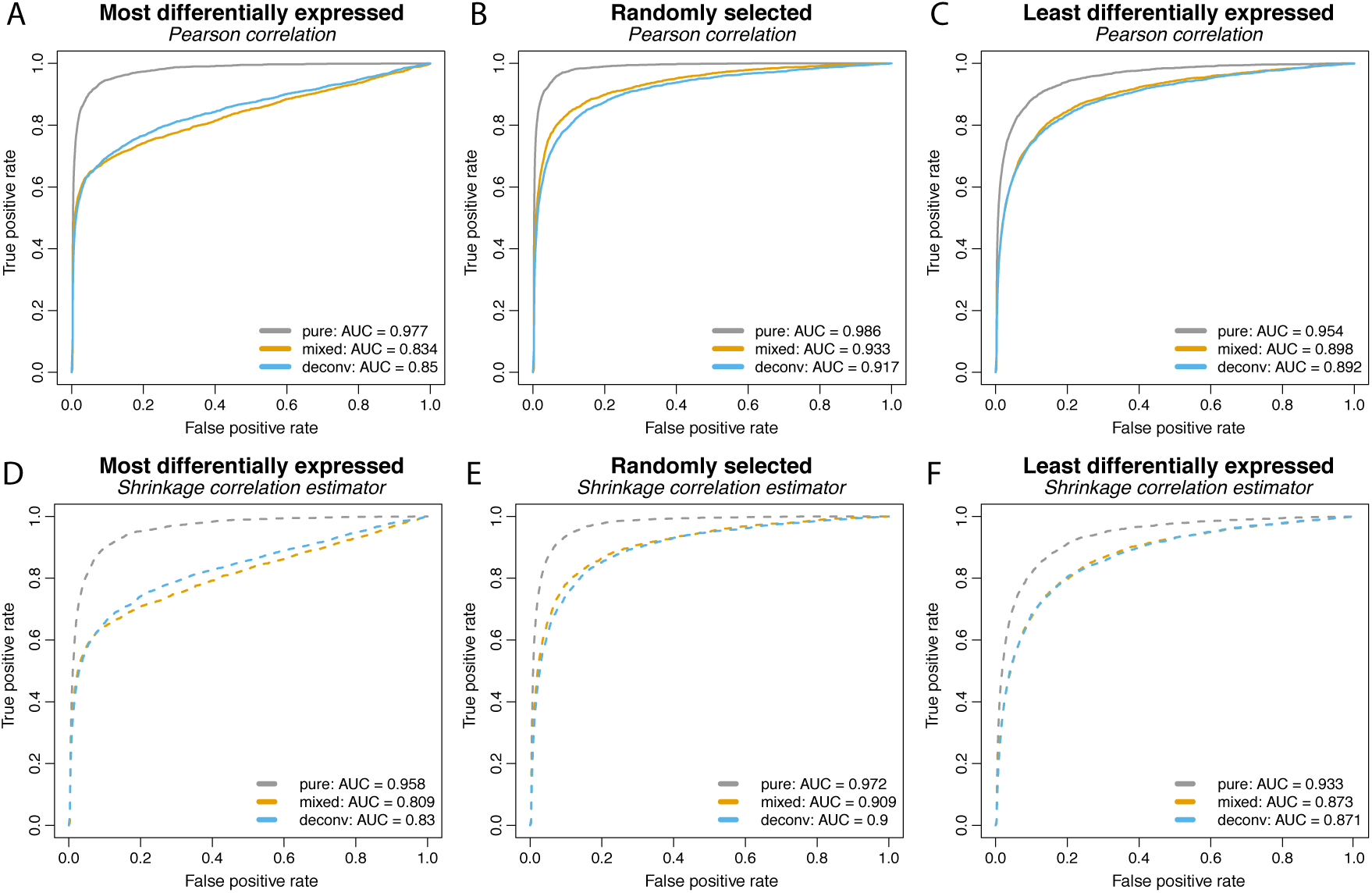
Counterpart of Figure 3 with proportions generated from the Beta(8,2), i.e. large proportions (on average 0.8) for signal-carrying cell-type in the mixture.

**Figure S5:**
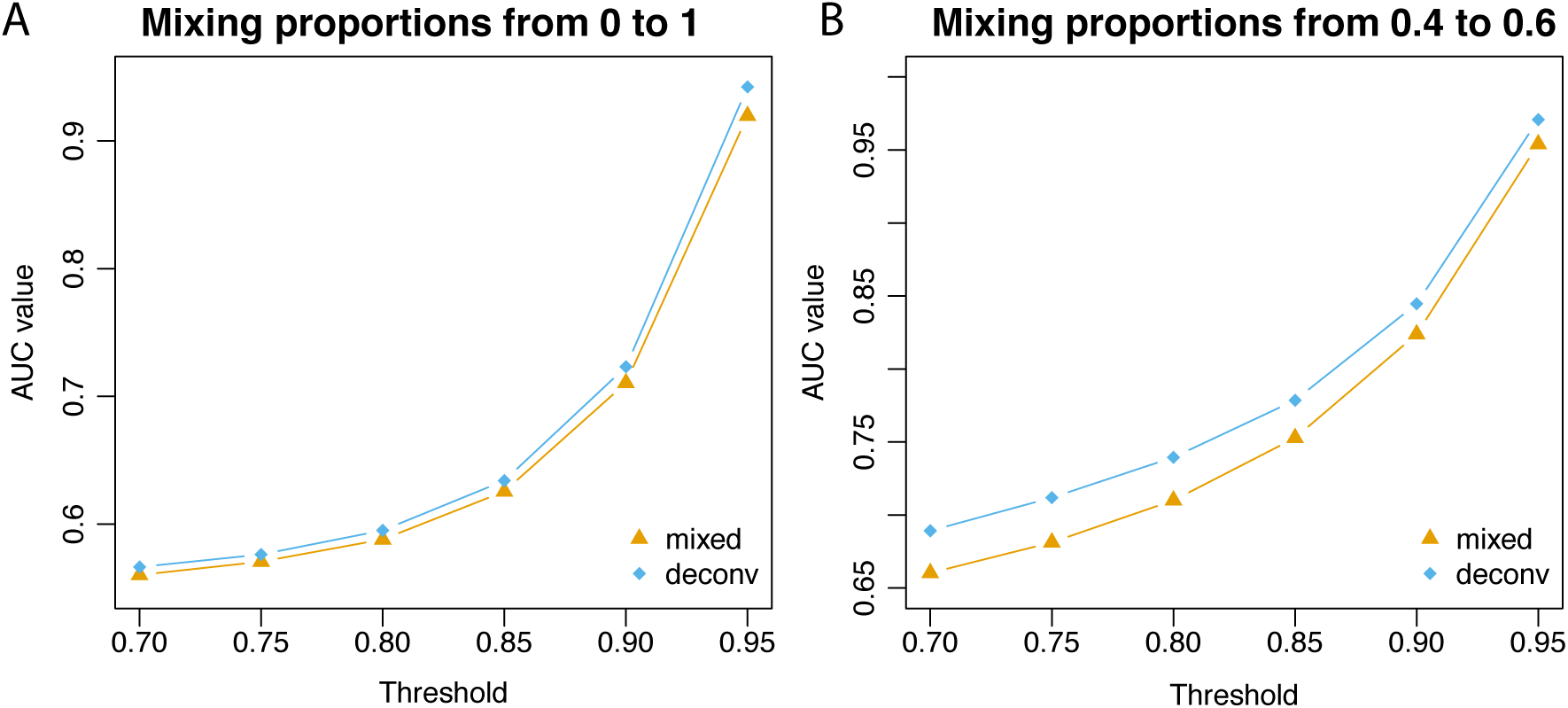
AUC values for `microRNAome` network estimation with varying thresholds.

